# β-cell-specific Non-invasive Ultrasound Stimulation to Enhance Insulin Release and Glucose Control in Mice

**DOI:** 10.1101/2025.09.03.673936

**Authors:** Yong Wu, Xinyi Zhao, Yizhou Jiang, Congmin Chen, Langzhou Liu, Xuandi Hou, Quanxiang Xian, Jinghui Guo, Lei Sun

**Affiliations:** Department of Biomedical Engineering, The Hong Kong Polytechnic University, Hung Hom 999077 Hong Kong SAR, PR China; School of Medicine, Chinese University of Hong Kong (Shenzhen), Shenzhen, China

**Keywords:** Ultrasound, Diabetes, Insulin Release, Mechanosensitive Ion channels, Microbubble

## Abstract

Diabetes poses a significant global health burden, with complications such as cardiovascular disease, stroke, and kidney failure. While insulin therapy is central to type 2 diabetes (T2D) management, its limitations—including rapid degradation and the need for frequent injections—highlight the demand for non-invasive alternatives. Here, we present an ultrasound (US)-mediated approach to enhance insulin release by selectively stimulating pancreatic β-cells via targeted microbubbles (MBs). *In vitro* experiments using RINm5F β-cells demonstrated that US-MB stimulation induces significant calcium influx and subsequent insulin release. In addition, this method effectively decreased blood glucose levels in mice by promoting insulin release. Mechanistic studies revealed that mechanosensitive ion channels play a pivotal role, as their inhibition (via GdCl_3_) abolished the ultrasonic effect. Importantly, the approach exhibited high biosafety, with no detectable cell death or tissue damage. Our findings establish ultrasound-stimulated β-cell targeting as a promising non-invasive strategy for diabetes treatment, offering a potential alternative to conventional insulin therapy.

## 1. Introduction

Diabetes is a chronic endocrine disorder characterized by persistent hyperglycemia, ranking among the most prevalent and rapidly growing global health crises [1]. Its severe complications—including vision loss, kidney failure, cardiovascular disease, stroke, and lower limb amputations—contribute to rising morbidity and mortality, posing a significant socioeconomic burden [2, 3]. Type 2 diabetes (T2D), accounting for 90–95% of cases, arises from insulin resistance and dysfunctional glucose-dependent insulin release [4]. While lifestyle interventions (e.g., exercise, weight management) can mitigate early-stage T2D [5], many patients, particularly the elderly, are unsuitable for such approaches [6]. Pharmacological therapies that enhance insulin release remain a clinical mainstay, yet their efficacy often wanes over time, and systemic side effects persist [7, 8]. Insulin therapy, though effective for glycemic control when oral medications fail [9], is hampered by peptide instability, rapid degradation, and the need for frequent injections—leading to hypoglycemia risks, patient discomfort, and logistical challenges in storage and administration [10].

To address these limitations, physical energy-based modalities—such as light [11], magnetic fields [12], and electrical stimulation [13]—have emerged as on-demand regulators of insulin release. While promising, these methods face translational hurdles, including poor spatiotemporal precision, limited tissue penetration, or cumbersome hardware. Thus, a pressing need remains for non-invasive, targeted strategies to restore insulin release and improve glucose management. Ultrasound (US) has recently gained attention as a potential solution, offering deep tissue penetration, high spatiotemporal precision, and excellent biosafety. In 2017, pioneering work demonstrated that US could stimulate insulin release from pancreatic β-cells *in vitro* [14]. However, critical gaps persist: (1) direct evidence of US-mediated glucose regulation *in vivo* remains scarce [15], and (2) off-target effects (e.g., glucagon elevation due to non-specific α-cell activation) may counteract therapeutic benefits [16]. For US to become a viable diabetes therapy, precise β-cell targeting is essential to minimize hormonal interference.

We hypothesize that targeted β-cell stimulation, achieved via microbubble (MB)-amplified ultrasound, can selectively enhance insulin release without perturbing glucagon release. By coupling MBs with β-cells, we aim to localize US-induced sonostimulation, triggering calcium influx and insulin exocytosis while avoiding off-target effects [17]. Here, we demonstrate that MB-enhanced US selectively stimulates RINm5F β-cells *in vitro*, inducing calcium-dependent insulin release, this effect is achieved by activating mechanosensitive channels. Crucially, this approach significantly lowers blood glucose in mice, providing the first direct evidence of US-mediated glycemic regulation *in vivo*. Our findings establish a foundation for non-invasive, stimulation-based therapies targeting pancreatic β-cells, offering a potential alternative to conventional diabetes management.

## 2. Results

### 1. Microbubble characterization and biosafety evaluation

Microbubbles (MBs) were synthesized via agitation of a heated lipid solution in the presence of air, followed by PEGylation to enhance stability and biocompatibility [18]. Size isolation yielded MBs with a uniform diameter distribution (average: 2.01 ± 0.31 µm; Fig. 1A, B), ensuring minimal risk of uncontrolled collapse or cell -damaging inertial cavitation during experiments. Zeta potential measurements confirmed colloidal stability (-18.1 ± 5.43 mV), consistent with PEG-coated MBs under physiological conditions.

**Fig. 1.**
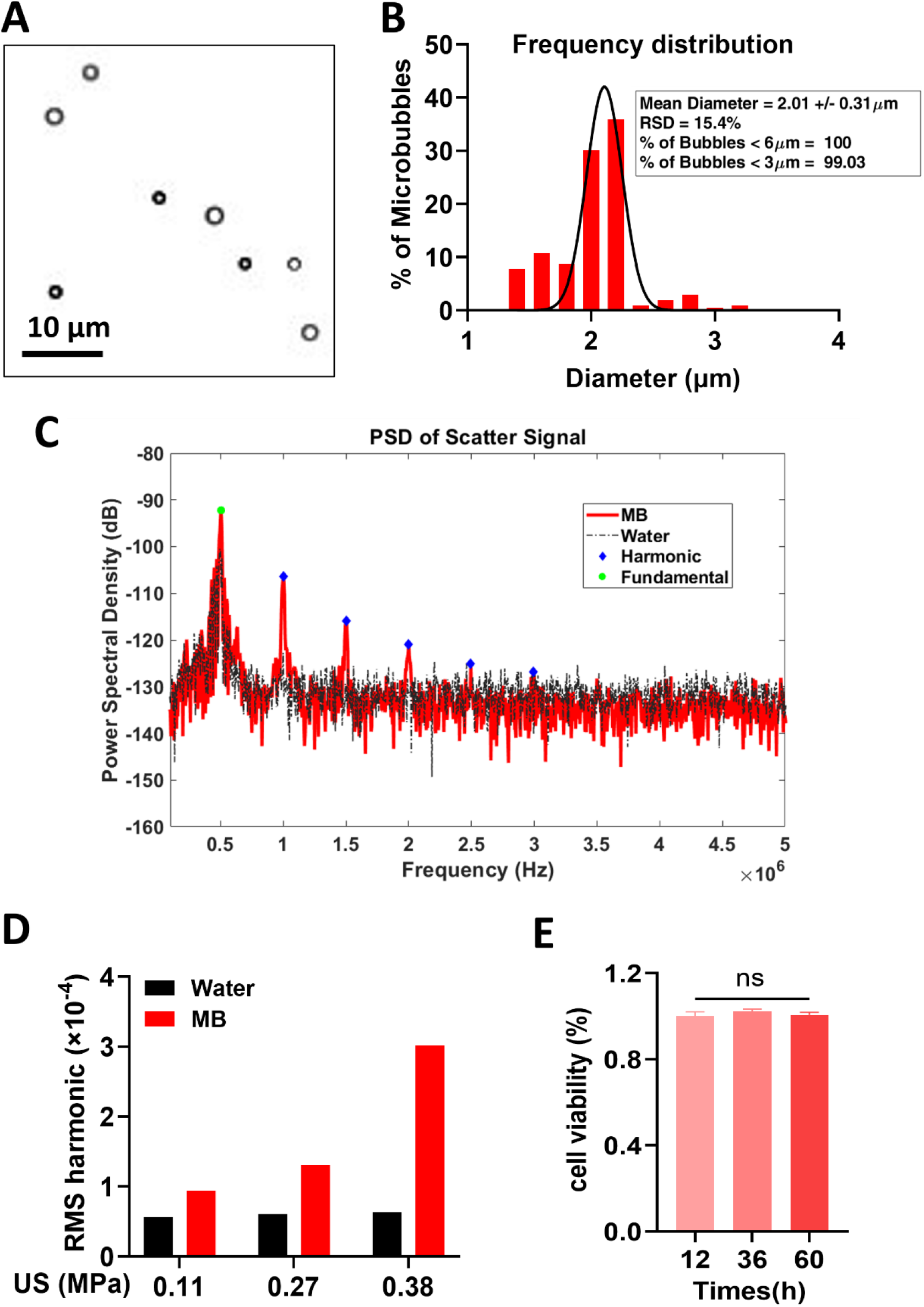
Characterization of microbubbles (MBs) and their interaction with ultrasound. **(A)** Bright-field microscopy image of synthesized MBs (scale bar: 10 µm). **(B)** Size distribution of MBs (mean diameter: 2.01 ± 0.31 µm), analyzed from (A) using MATLAB. **(C)** Frequency spectrum of backscattered signals from MB suspension during sonication (0.5 MHz, 50-cycle tone burst, 0.38 MPa peak negative pressure). Fundamental frequency (green dot) and harmonics (blue diamonds) are labeled. **(D)** RMS power of 2nd–8th harmonics in degassed water (black) versus MB suspension (red) across increasing acoustic pressures (0.11–0.38 MPa). **(E)** Cytocompatibility of MBs assessed by MTS assay. RINm5F cell viability (normalized to MB-free controls) after 12–60 hrs exposure to MBs (n = 3 biological replicates; mean ± SEM; ns: not significant by one-way ANOVA).

To quantify MB-mediated ultrasound amplification, we analyzed acoustic signals using a passive cavitation detector. Nonlinear harmonic scattering—a hallmark of stable cavitation—was assessed via root mean square (RMS) power within 40 kHz spectral windows centered at harmonic frequencies (Fig. 1C). In pure water, harmonic power remained low and intensity-independent, reflecting the absence of cavitation nuclei. In contrast, MB suspensions exhibited intensity-dependent harmonic amplification (Fig. 1D), confirming that MBs efficiently convert ultrasound energy into localized mechanical stimuli at low intensities (0.11–0.38 MPa).

To evaluate the biosafety of MBs, RINm5F β-cells co-cultured with MBs for 72 hrs showed no reduction in proliferation activity (MTS assay; Fig. 1E), demonstrating excellent biocompatibility. This result validates the safety of MBs for *in vitro* and *in vivo* applications.

### 2. Ultrasound-induced Ca^2+^ influx and insulin release

To evaluate ultrasound-triggered β-cell activation, we performed live-cell Ca^2+^ imaging in RINm5F cells loaded with Fura-2 AM (Fig. 2A). Cells were exposed to ultrasound (0.5 MHz, 0.5 ms pulse width, 1 ms interval, 300 ms duration, 3 s repetition) in the presence of microbubbles (MBs; 0–1.65 × 10^9^/mL) (Fig. 2B, Extended data 1). At 0.14 MPa, ultrasound induced MB concentration-dependent Ca^2+^ influx, with saturation observed at 1.65 × 10^9^ MBs/mL (Fig. 2C, D, Extended data 2). We next assessed the acoustic pressure threshold for Ca^2+^ mobilization (0.07–0.38 MPa). Robust, dose-dependent Ca^2+^ responses occurred only in MB-treated cells (p < 0.001 vs. MB-free controls at all pressures; Fig. 2E), confirming MBs are essential for ultrasound-mediated stimulation at low intensities.

**Fig. 2.**
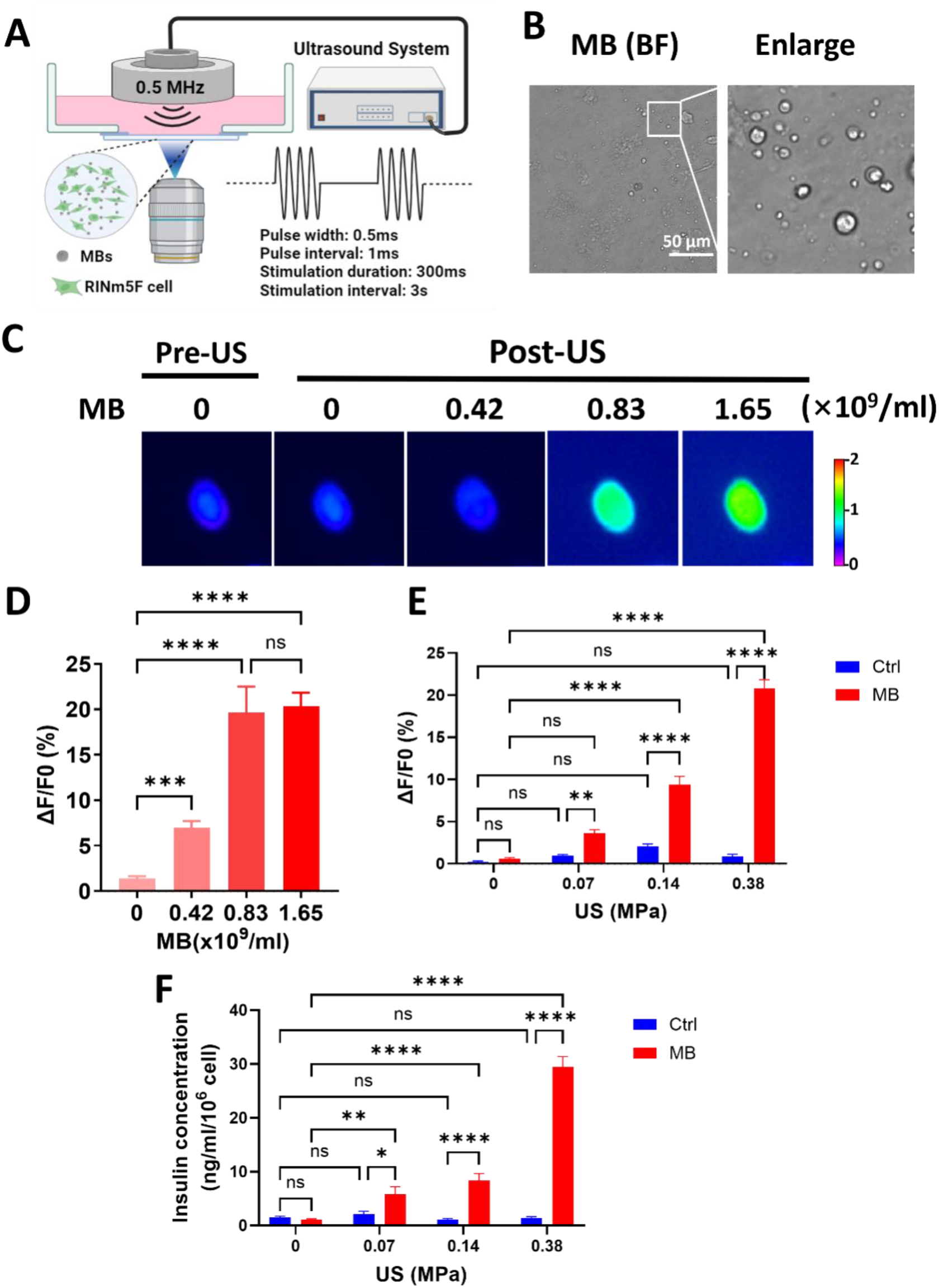
Microbubble-enhanced ultrasound stimulation triggers Ca^2+^ influx and insulin release in RINm5F β-cells. **(A)** Experimental setup for ultrasound stimulation (0.5 MHz, 0.5 ms pulse width, 1 ms interval, 300 ms duration, 3 s repetition) and live-cell Ca^2+^ imaging. MBs were suspended in culture medium above adherent RINm5F cells. **(B)** Bright-field microscopy showing uniform MB distribution among cells (scale bar: 50 μm). **(C)** Representative Fura-2 fluorescence images showing intracellular Ca^2+^ levels before and after ultrasound stimulation (0.14 MPa) with MBs (0-1.65×10^9^/mL). **(D)** Ca^2+^ response kinetics under 0.14 MPa ultrasound with increasing MB concentrations (n = 28-30 cells/group; mean ± SEM; ***p < 0.001, ****p < 0.0001, one-way ANOVA with Dunn’s post-hoc tests). **(E)** Ultrasound intensity-dependent Ca^2+^ responses with/without MBs (1.65×10^9^/mL; n = 10-55 cells/group). **(F)** Insulin release measured by ELISA 15 min post-stimulation (n = 4 biological replicates). Data: mean ± SEM; *p < 0.05, **p < 0.01, ***p < 0.001, ****p < 0.0001 (two-way ANOVA).

As Ca^2+^ influx triggers insulin exocytosis [19], we quantified released insulin via ELISA. Ultrasound + MBs elicited significant, intensity-dependent insulin release (p < 0.001 at 0.14 and 0.38 MPa), while no response occurred without MBs (Fig. 2F). This demonstrates that MB-enhanced ultrasound selectively activates β-cells without off-target effects.

### 3. Microbubble-enhanced ultrasound stimulation promotes insulin release and improves glycemic control *in vivo*

To evaluate the therapeutic potential of MB-enhanced ultrasound stimulation, we subcutaneously implanted RINm5F cells embedded in Matrigel into nude mice (Fig. 3A). Following a 6-hour fast, mice received the ultrasound treatment under isoflurane anaesthesia. Specifically, MBs were first administered to the cell transplantation site, followed by an intraperitoneal injection of glucose (2 g/kg). Ultrasound stimulation (0.38 MPa, 15 min duration, parameters consistent with prior protocols; see Fig. 2A) was immediately applied to the same region, with acoustic coupling gel minimizing signal attenuation (Fig. 3A).

**Fig. 3.**
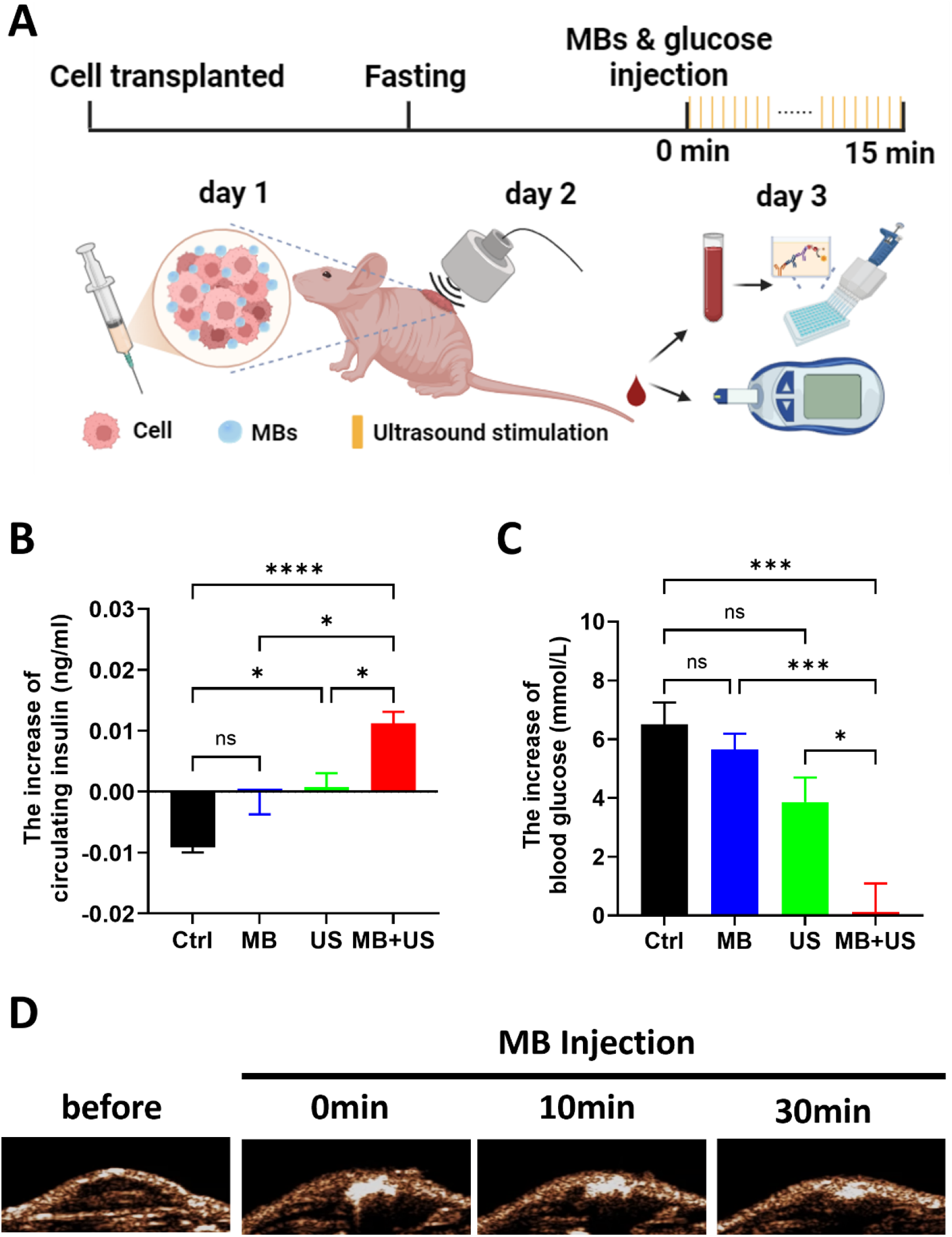
Microbubble-enhanced ultrasound stimulation promotes insulin release and improves glycemic control in mice. **(A)** Experimental timeline and setup: RINm5F cells were implanted subcutaneously 24 hrs before the intraperitoneal glucose injection (2 g/kg). The MB+US group received microbubble injection followed by ultrasound stimulation (0.5 MHz, 0.38 MPa, 50% duty cycle, 300 ms pulse duration, 3 s interval) for 15 min post-glucose challenge. Blood was collected for glucose and insulin measurements at 0 (before US) and 15 min (immediately after US). **(B)** Plasma insulin levels normalized to baseline (t=0). MB+US group showed significant enhancement vs. Ctrl and US-only groups (n=5 mice/group; mean ± SEM; *p<0.05, ****p<0.0001, one-way ANOVA with post-hoc Tukey’s test). **(C)** Blood glucose performance normalized to baseline (t=0). MB+US group exhibited accelerated glucose clearance (n=5 mice/group; mean ± SEM; *p<0.05, ***p<0.001 vs. Ctrl at matched timepoint, one-way ANOVA with post-hoc Tukey’s test). **(D)** Representative nonlinear contrast ultrasound images confirming stable MB distribution at implantation sites.

As expected, glucose challenge rapidly elevated blood glucose levels in all groups (Fig. 3C). While control animals showed decreased insulin response (likely due to isoflurane affect [20, 21]), the MB+US group exhibited significantly higher insulin levels compared to controls (p < 0.0001; Fig. 3B). Ultrasound alone (US group) induced only marginal insulin elevation (Fig. 3B), underscoring the essential role of MBs in amplifying the bioeffects of ultrasound stimulation.

The enhanced insulin release in MB+US mice translated to superior glucose maintenance, with significantly lower blood glucose levels at 15 min post-challenge versus controls (p < 0.001; Fig. 3C). Notably, MBs remained stable throughout the experiment, with ultrasound imaging confirming persistent contrast signals 30 min post-injection (Fig. 3D), indicating sustained cavitation activity. These results demonstrate that MB-localized ultrasound stimulation of transplanted β-cells can effectively augment glucose-dependent insulin release and improve glycemic control in vivo, offering a potential strategy for non-invasive diabetes management.

### 4. Mechanistic insights into microbubble-enhanced insulin release

Since Ca^2+^ influx directly triggers insulin exocytosis [19], we hypothesized that MB-enhanced ultrasound activates mechanosensitive ion channels (Fig. 4A). qPCR confirmed robust expression of multiple mechanosensitive channels in RINm5F cells, with selective subtypes exhibiting particularly high abundance (Fig. 4B). To dissect the mechanism, we used gadolinium chloride (GdCl_3_), a broad-spectrum mechanosensitive channel blocker [22]. At 0.14 MPa (parameters matching Fig. 2A), GdCl_3_ abolished ultrasound-induced Ca^2+^ influx (Fig. 4C, D), indicating channel activation dominates. Consistent with Ca^2+^ data, GdCl_3_ reduced insulin release by 75% under 0.14 MPa US+MB stimulation (Fig. 4E). The residual insulin release in GdCl_3_-treated cells (statistically insignificant vs. controls) likely reflects slight membrane perturbation. These results demonstrate that MB-enhanced ultrasound primarily stimulates insulin release via mechanosensitive channel activation.

**Fig. 4.**
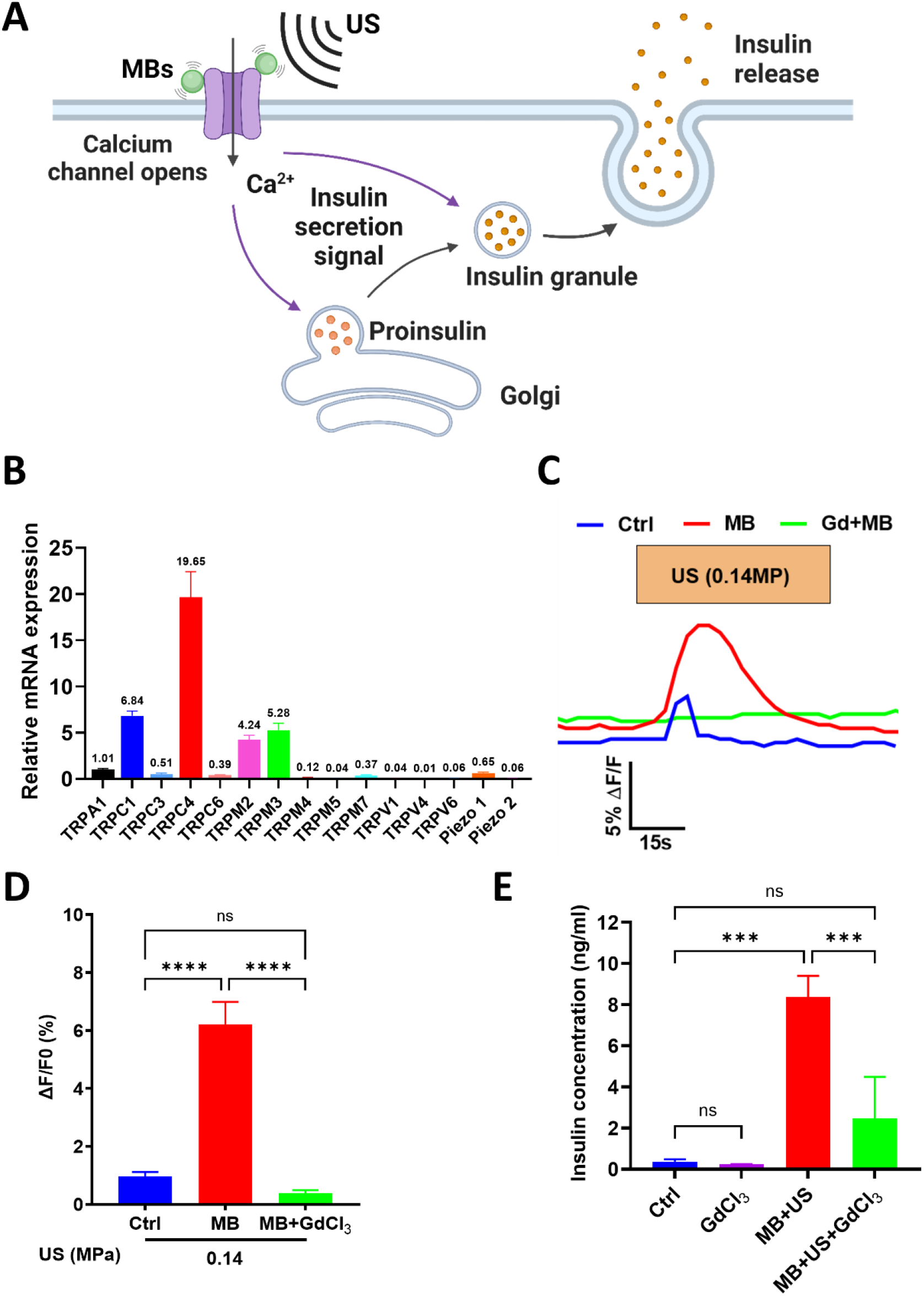
Mechanistic role of mechanosensitive ion channels in microbubble-enhanced ultrasound stimulation of insulin release. **(A)** Proposed mechanism: MB-mediated ultrasound activates mechanosensitive channels to induce Ca^2+^ influx and insulin exocytosis. **(B)** qPCR analysis of mechanosensitive channel expression in RINm5F cells (n=3 biological replicates). **(C)** Representative Fura-2 traces showing Ca^2+^ dynamics during 0.14 MPa ultrasound stimulation in control, MB-only, and MB+GdCl_3_ (50 μM) groups. **(D)** Quantified Ca^2+^ responses of (C) (n=31-57 cells/group; mean ± SEM; ****p<0.0001, one-way ANOVA with post-hoc Tukey’s test)). **(E)** Insulin release after 0.14 MPa stimulation (n=3 biological replicates; mean ± SEM; ***p<0.001, one-way ANOVA with post-hoc Tukey’s test).

### 5. Biosafety evaluation of MB-enhanced ultrasound stimulation

Given the potential for cavitation-induced cellular effects, we conducted comprehensive biosafety assessments of our MB-enhanced ultrasound approach. Initial *in vitro* studies using propidium iodide (PI) staining confirmed excellent cellular viability across all treatment groups, with no significant cell death observed following ultrasound stimulation at any intensity (Fig. 5A). Further investigation of transient membrane permeability revealed minimal reversible sonoporation, as evidenced by PI-positive rates below 0.5% at therapeutic intensities (0.07-0.14 MPa) and only marginally higher rates (∼1.1%) at 0.38 MPa (Fig. 5B). These findings corroborate our mechanistic studies, demonstrating that insulin release at lower intensities primarily occurs through bio-safe mechanosensitive channel activation rather than membrane disruption.

**Fig. 5.**
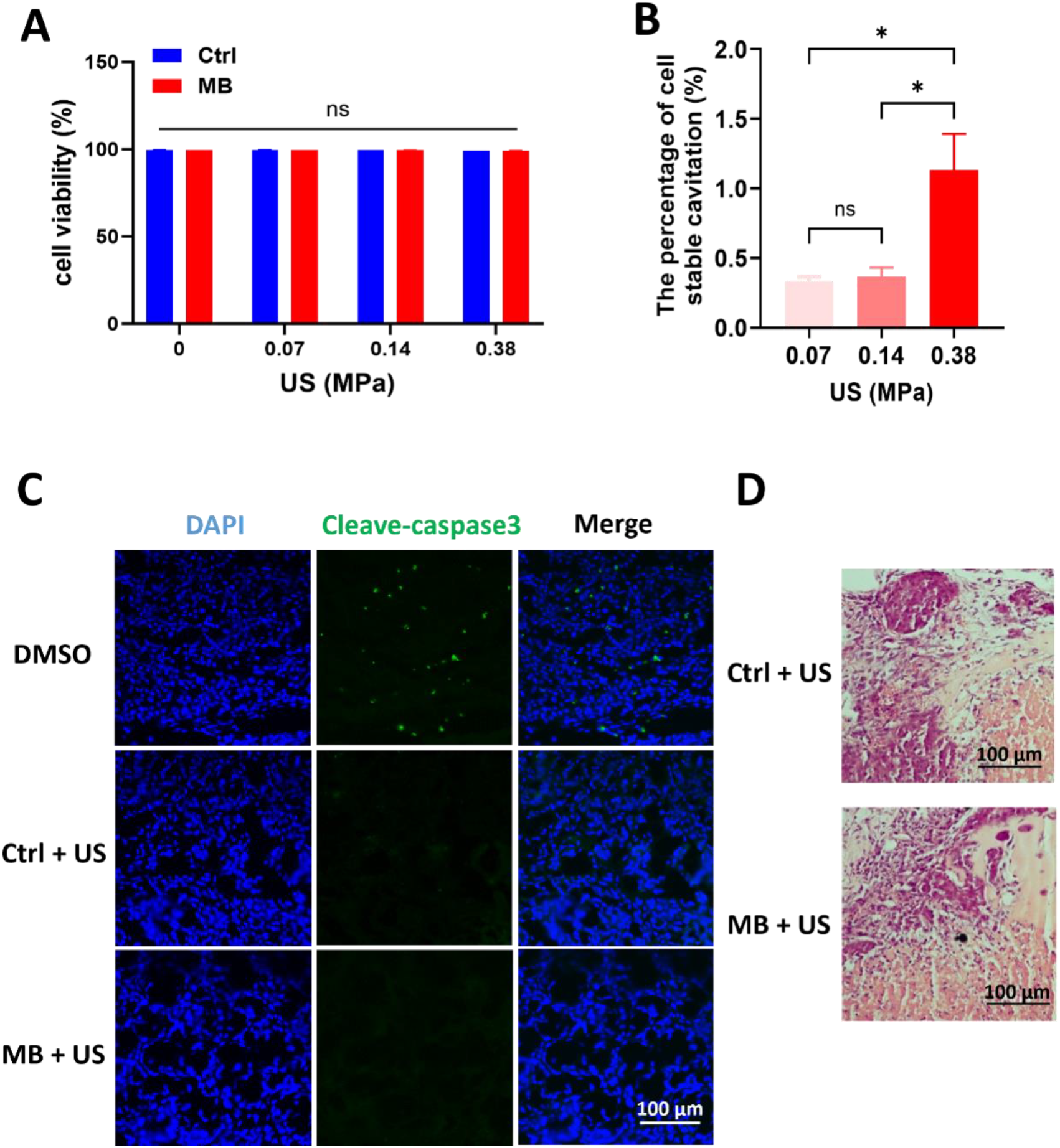
Biosafety assessment of microbubble-enhanced ultrasound stimulation. **(A)** Cell viability following ultrasound treatments (conditions matching Fig. 2E, F), assessed by propidium iodide (PI) exclusion (n=3 biological replicates; ns: not significant by two-way ANOVA). **(B)** Sonoporation quantification via PI uptake during treatment (n=3; mean ± SEM; *p<0.05 vs control, one-way ANOVA with Tukey’s test). **(C)** Cleaved caspase-3 immunofluorescence in transplanted tissues (DMSO: positive control; scale bar: 100 μm). **(D)** H&E staining of treated tissues showing preserved morphology (scale bar: 100 μm).

For *in vivo* safety validation, we examined transplanted tissues from treated mice through two complementary approaches. Cleaved caspase-3 immunofluorescence staining, a sensitive apoptosis marker, showed no detectable signals in either ultrasound-treated or MB+US groups, in contrast to clear positive signals in DMSO-treated controls (Fig. 5C). Consistent with these results, hematoxylin and eosin (H&E) staining revealed preserved tissue architecture with no evidence of structural damage (Fig. 5D).

Collectively, these results establish the strong biosafety profile of MB-enhanced ultrasound stimulation. The approach maintains cellular integrity at therapeutic intensities while achieving effective insulin release through controlled sonostimulation, with only minimal and reversible membrane effects observed at higher pressures. Both *in vitro* and *in vivo* assessments confirm the absence of significant cytotoxicity, apoptosis, or tissue damage, supporting the potential clinical translatability of this technique for diabetes management.

## 3. Discussion

This study establishes the first proof-of-concept for blood glucose regulation through microbubble-assisted, β-cell-specific ultrasound stimulation in murine models. Our findings reveal that localized ultrasound application with microbubble (MB) mediation significantly enhances insulin release from transplanted RINm5F β-cells, effectively delaying glucose elevation in glucose-challenged animals. Through systematic optimization of ultrasound parameters and MB concentrations, we developed a dose-dependent insulin modulation paradigm while maintaining biosafety, as evidenced by reversible membrane sonoporation and absence of histopathological cellular damage. These results provide novel mechanistic insights into ultrasound-mediated glycemic control in mammals.

Existing evidence supports ultrasound-modulated insulin release in pancreatic β-cells [14-16], laying the foundation and hope for ultrasound-based diabetes therapy. Nonetheless, the development of specific therapeutic methods and significant efficacy in diabetic mouse model remains to be demonstrated. Prior investigations have primarily employed non-specific pancreatic ultrasound stimulation. Crucially, the pancreas contains multiple endocrine cell populations - including insulin-antagonizing glucagon-secreting islet α-cells and acinar cells (constituting the pancreatic cellular majority) - that reside in intimate spatial association with islet β-cells [23]. This anatomical proximity renders conventional ultrasound approaches incapable of cellular discrimination, potentially inducing counterproductive α-cell activation that obscures glucose-regulatory outcomes. This suggests that β-cell-specific targeting represents the critical determinant for achieving precise ultrasound-mediated glycemic control. The pivotal advancement of this study lies in demonstrating that ultrasound-targeted β-cell activation not only enhances insulin release but also effectively mitigates acute hyperglycemic spikes *in vivo*. This achievement validates the feasibility of selective β-cell modulation for glucose management and provides direct experimental evidence supporting the potential of ultrasound-based strategies for future diabetes intervention.

While our transplantation model and MB-based system confirm the principle of ultrasound-specific β-cell regulation, clinical translation remains constrained by inherent MB limitations (e.g., instability and systemic delivery challenges). Further strategy that can specifically enhance the sonosensitivity of β-cell with long lifetime may achieve a non-invasive, on-demand ultrasound-mediated glycemic control paradigm to reshape diabetes therapy.

## Acknowledgement

This work was financially supported by the Hong Kong Research Grants Council Collaborative Research Fund (C5053-22 GF), General Research Fund (15126524 and 15224323), National Key Research and Development Program of Ministry of Science and Technology of China (2023YFC2410900), and internal funding from the Hong Kong Polytechnic University (G-SACD), Research Center for Non-invasive Brain Computer Interface (1-CE0M), and Research Institute of Smart Ageing (1-CDJM). The authors would like to thank the facility and technical support from the University Research Facility in Life Sciences (ULS) and University Research Facility in Behavioral and Systems Neuroscience (UBSN) of The Hong Kong Polytechnic University.

## Author Contributions

Conceptualization, Y.W., X.Z., J.G., and L.S.; Methodology, Y.W., X.Z., Y.J., X.H., Q.X., J.G., and L.S.; Investigation, Y.W., X.Z., and J.G.; Formal Analysis, Y.W., X.Z., and Y.J.; Writing—Original Draft, Y.W., X.Z., and L.S.; Writing—Review and Editing, Y.W., X.Z., J.G., Y.J., C.C., L.L., and L.S.; Supervision, J.G. and L.S.; Funding Acquisition, L.S.

## 4. Methods and Materials

### 4.1. Cell culture

Rat beta-insulinoma cells (RINm5F) were obtained from ATCC and cultured in RPMI 1640 medium supplemented with 10% fetal bovine serum (FBS) and 1% penicillin-streptomycin (all from Gibco). The cells were maintained in a humidified incubator at 37°C with 5% CO_2_. For Ca^2+^ imaging with direct ultrasound treatment, cells were seeded in 35-mm confocal dishes at a density of 1×10^6 cells/dish, cultured overnight, and subjected to ultrasound treatment the following day.

### 4.2. Cell MTS

The MTS assay was performed according to the manufacturer’s instructions (ab197010). In brief, 8000 RINm5F cells were cultured in the 96-well-plate. Cells were cultured under standard conditions or co-cultured with microbubbles (MBs) under identical conditions as the control group. MTS reagent was added to the cell medium at the 12h, 36h, and 60h time points. Absorbance was measured using a microplate reader (Labexim LEDETECT 96), and cell numbers were subsequently calculated. Cell viability was calculated as “the cell number of MBs group” / “the cell number of control group” *100% at each time point.

### 4.3. Ultrasound stimulation system and protocol

The ultrasound stimulation system consisted of a function generator and a commercial transducer (I7-0012-P-SU, Olympus) to produce 200 tone burst pulses at a center frequency of 500 kHz and a repetition frequency of 1 kHz with a duty cycle of 50%. The output intensity was maintained between 0.1 and 0.5 MPa, with a 3 second interval between pulses. During the Ca^2+^ imaging measurement, the transducer was tilted at 90° to the confocal dish. Temperature was monitored following ultrasound treatment using the same parameters as in our previous studies, with no significant temperature increase observed (data not shown).

### 4.4. Ca^2+^ Imaging

Cells seeded in 35mm confocal dishes were washed with the bath solution containing (in mM): NaCl 130, KCl 5, MgCl_2_ 1, CaCl_2_ 2.5, Glucose 10, HEPES 20 (pH 7.4),then incubated with 3 µM Fura-2 (F1200, Thermo Fisher Scientific Inc, MA, USA) in bath solution at 37 °C for 30 min, Subsequently, the dishes were transferred to a fluorescence microscope (Eclipse Ti; Nikon, Tokyo, Japan) for intracellular Ca^2+^ measurements. Fluorescence was alternatively excited by dual-wavelength 340 and 380 nm with an interval of 3 s, and the emitted fluorescent lights were collected at 510 nm. Selected cells were incubated with 20 µM GdCl_3_ dissolved in water. The drug was still maintained in the bath solution throughout Ca^2+^ imaging measurement.

### 4.5. Mice blood glucose and insulin measurement

Mice were obtained from Centralized Animal Facilities (CAF) at Hong Kong Polytechnic University or the Laboratory Animal Services Centre at The Chinese University of Hong Kong. All animals were housed at CAF and experimentally procedures were approved by the Animal Subjects Ethics Sub-committee at Hong Kong Polytechnic University (22-344) in DH/HT&A/8/2/4 Pt.12). Mice were fasted for 6 h prior to intraperitoneal glucose injection (2 g/kg body weight). Blood was collected via the tail before and after 15 minutes of glucose injection. Glucose and insulin levels in the collected blood were determined by glucose test strip (Bayer HealthCare LLC) and insulin ELISA kit (ImmunoDiagonstics 32380), respectively. Ultrasound stimulation was delivered to the cell-transplanted region for 15 min following glucose injection.

### 4.6. Insulin ELISA

RINm5F cells were grown on 24-well plates. The cells were starved for 2 hours using glucose and FBS-free 1640 medium. MB that dissolved in PBS was added to the medium. Culture mediums were collected 15 minutes after the 10 mM glucose challenge. Insulin in the culture media was measured by ELISA following the manual of the manufacturer (ImmunoDiagonstics 33100).

### 4.7. qPCR Analysis

RNA extraction, reverse transcription, and real-time qPCR were performed as described in our previous study [24]. For each group, 1 µg of RNA was reverse transcribed. For real-time qPCR, 1 µL of cDNA from plasmid-transfected RINm5F cells was added to each qPCR reaction. Results are presented as fold changes relative to controls, expressed as mean ± SEM from three independent experiments. Primer sequences were as follows:

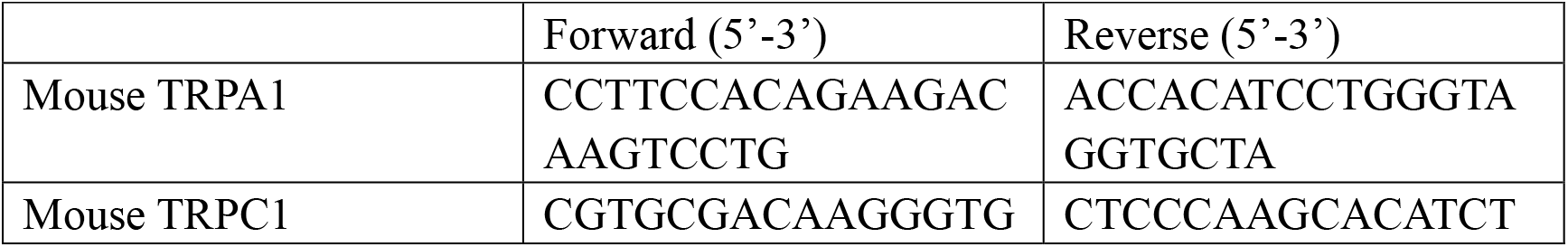
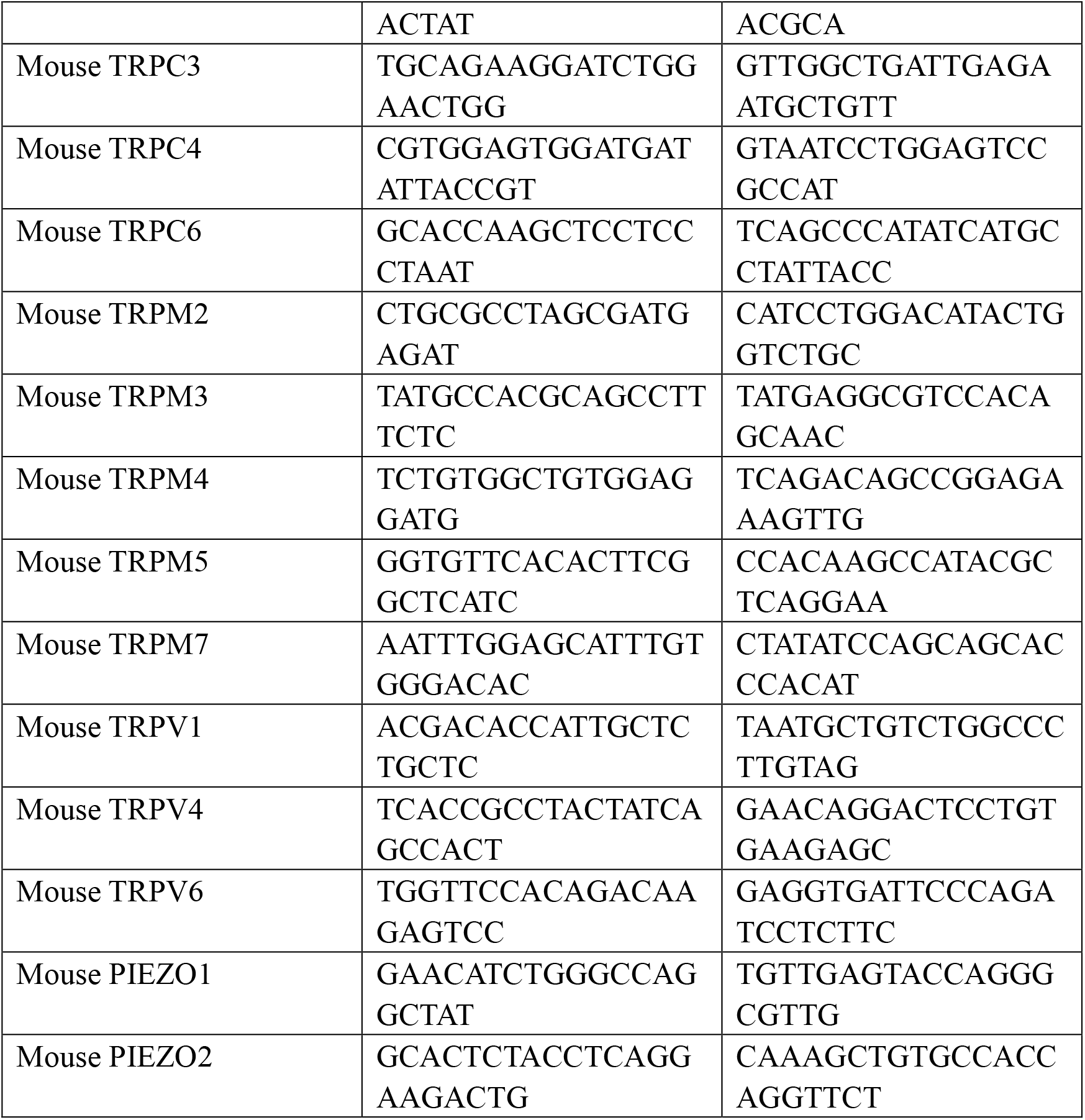

### 4.8. Immunofluorescence and HE staining

Fresh tumor tissues were excised and fixed in 4% formaldehyde, then permeabilized and blocked with a solution containing 0.1% Triton X-100, 5% normal goat serum, and 1% BSA in PBS for 1 hour at room temperature. Following buffer removal, samples were incubated overnight with primary antibodies diluted in the same buffer.Secondary antibody incubation was conducted the following day for 1 h at room temperature using antibodies diluted in the same buffer. After three washes, samples were dried and mounted using ProLong Glass Antifade Mountant with NucBlue (Life Technologies), covered with coverslips, and imaged with a confocal laser scanning microscope (TCS SP8; Leica). All steps from the secondary antibody incubation onwards were performed in the dark.

Primary antibodies included cleaved caspase-3 (9661T; Cell Signaling Technology) used at 1:500 dilution. Secondary antibodies (goat anti-rabbit IgG (H+L), Alexa Fluor 488; A11008, Invitrogen) were used at 1:1000 dilution.

The fixed tissues were also dehydrated using alcohol and vitrified in dimethylbenzene. Samples were embedded in paraffin, sectioned, and stained with hematoxylin and eosin (SBT10001; Sunteambio Biotechnology, Shanghai, China).

### 4.9. Statistical analysis

A minimum of 3 independent experiments were performed for all experiments shown. By this, we mean at least 3 separate ‘rounds’ of cell preparations, transfections, or RINm5F cell harvests used for various experiments. Wherever possible, multiple plates from each round were evaluated. The data were collected into GraphPad Prism sheets for statistical analysis and graph preparation. The normality of the data was assessed using the Shapiro-Wilk normality test. Next, the homogeneity of variances was evaluated using either the Brown-Forsythe test or Bartlett’s test. For data that were normally distributed and had homogeneous variances, ordinary one-way ANOVA was performed, followed by Tukey’s post-hoc test for multiple comparisons. If the data exhibited heterogeneity of variance, Welch’s ANOVA was used, followed by Dunnett’s T3 post-hoc test for multiple comparisons. Nonparametric data were analyzed using the Kruskal-Wallis test, followed by Dunn’s post-hoc test for multiple comparisons. A significance level of *p* values below 0.05 was considered statistically significant.

**Extended data 1:**
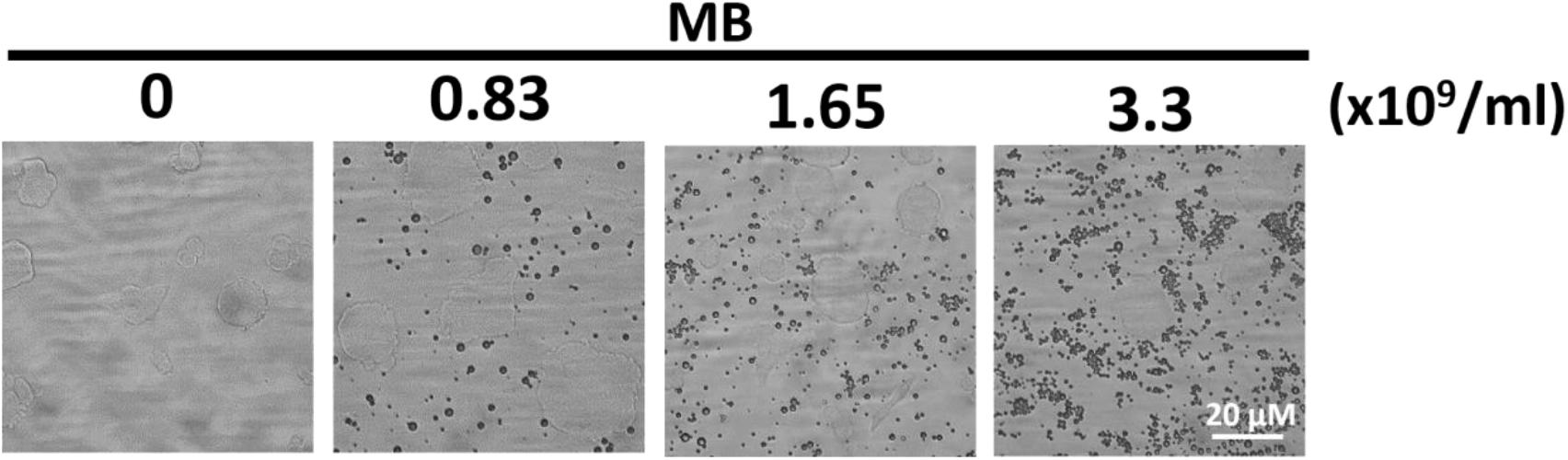
Schematic representation of different concentrations of microbubbles and cells were mixed in the white field.

**Extended data:**
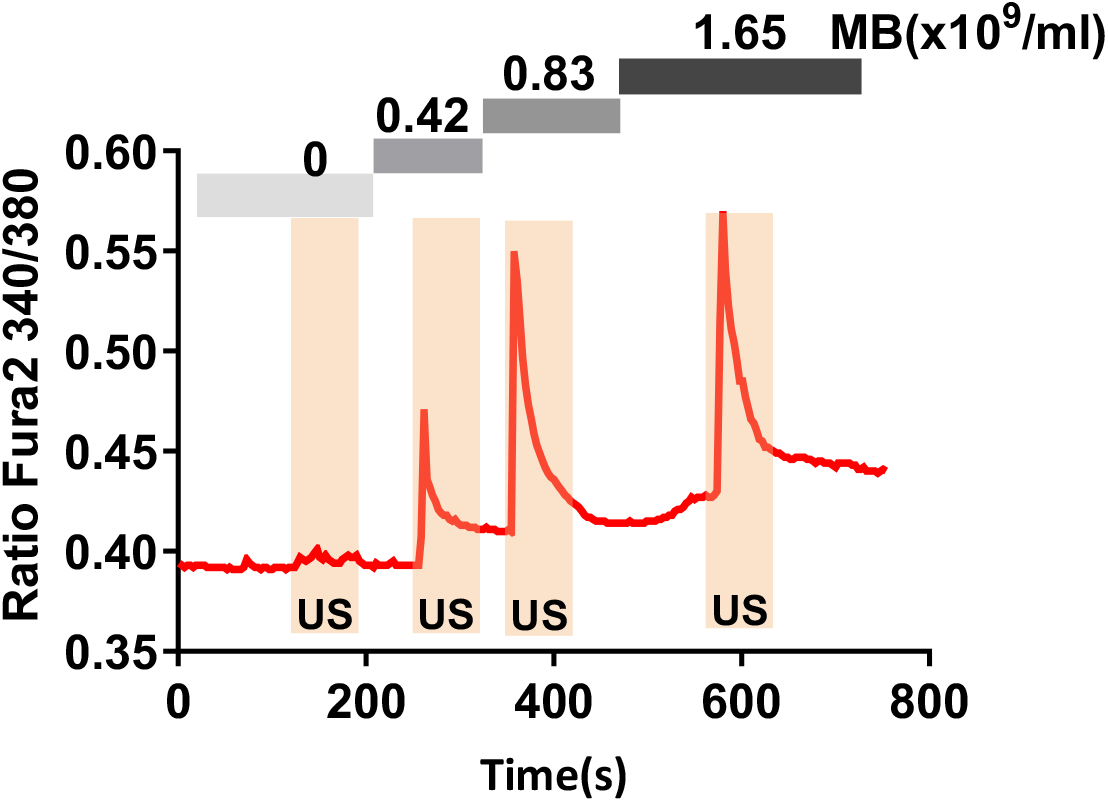
Time-course traces of Ca2+ measurement in the cells treated with different densities of microbubble under the same amplitude of ultrasound stimulation (0.07MPa).

